# Recognition of natural objects in the archerfish

**DOI:** 10.1101/2021.08.30.458062

**Authors:** Svetlana Volotsky, Ohad Ben-Shahar, Opher Donchin, Ronen Segev

## Abstract

Recognition of individual objects and their categorization is a complex computational task. Nevertheless, visual systems are able to perform this task in a rapid and accurate manner. Humans and other animals can efficiently recognize objects despite countless variations in their projection on the retina due to different viewing angles, distance, illumination conditions, and other parameters. Numerous studies conducted in mammals have associated the recognition process with cortical activity. Although the ability to recognize objects is not limited to mammals and has been well-documented in other vertebrates that lack a cortex, the mechanism remains elusive. To address this gap, we explored object recognition in the archerfish, which lack a fully developed cortex. Archerfish hunt by shooting a jet of water at aerial targets. We leveraged this unique skill to monitor visual behavior in archerfish by presenting fish with a set of images on a computer screen above the water tank and observing the behavioral response. This methodology served to characterize the ability of the archerfish to perform ecologically relevant recognition of natural objects. We found that archerfish can recognize an individual object presented under different conditions and that they can also categorize novel objects into known categories. Manipulating features of these objects revealed that the fish were more sensitive to object contours than texture and that a small number of features was sufficient for categorization. Our findings suggest the existence of a complex visual process in the archerfish visual system that enables object recognition and categorization.

## Introduction

For their survival, many animal species require the computational capacity to perform a range of complex object recognition tasks, from identifying a conspecific to recognizing a camouflaged predator, to classifying an item as edible (DiCarlo et al., 2012; Santos et al., 2001; Suboski and Templeton, 1989). Object recognition is defined as the ability to rapidly and accurately identify a specific object (Fig. 1A) or categorize objects into classes (Fig. 1B) despite substantial differences in the retinal representation of the object or across category members (DiCarlo et al., 2012). Variations in an object’s retinal image are typically caused by the different conditions under which the object is viewed; for example, its illumination, the viewing distance and angle, and other environmental characteristics (Biederman and Bar, 1999; DiCarlo and Cox, 2007; DiCarlo et al., 2012).The ability of animal brains to recognize objects in an efficient and accurate manner depends on powerful neural computations that enable classification and identification.

**Fig. 1.**
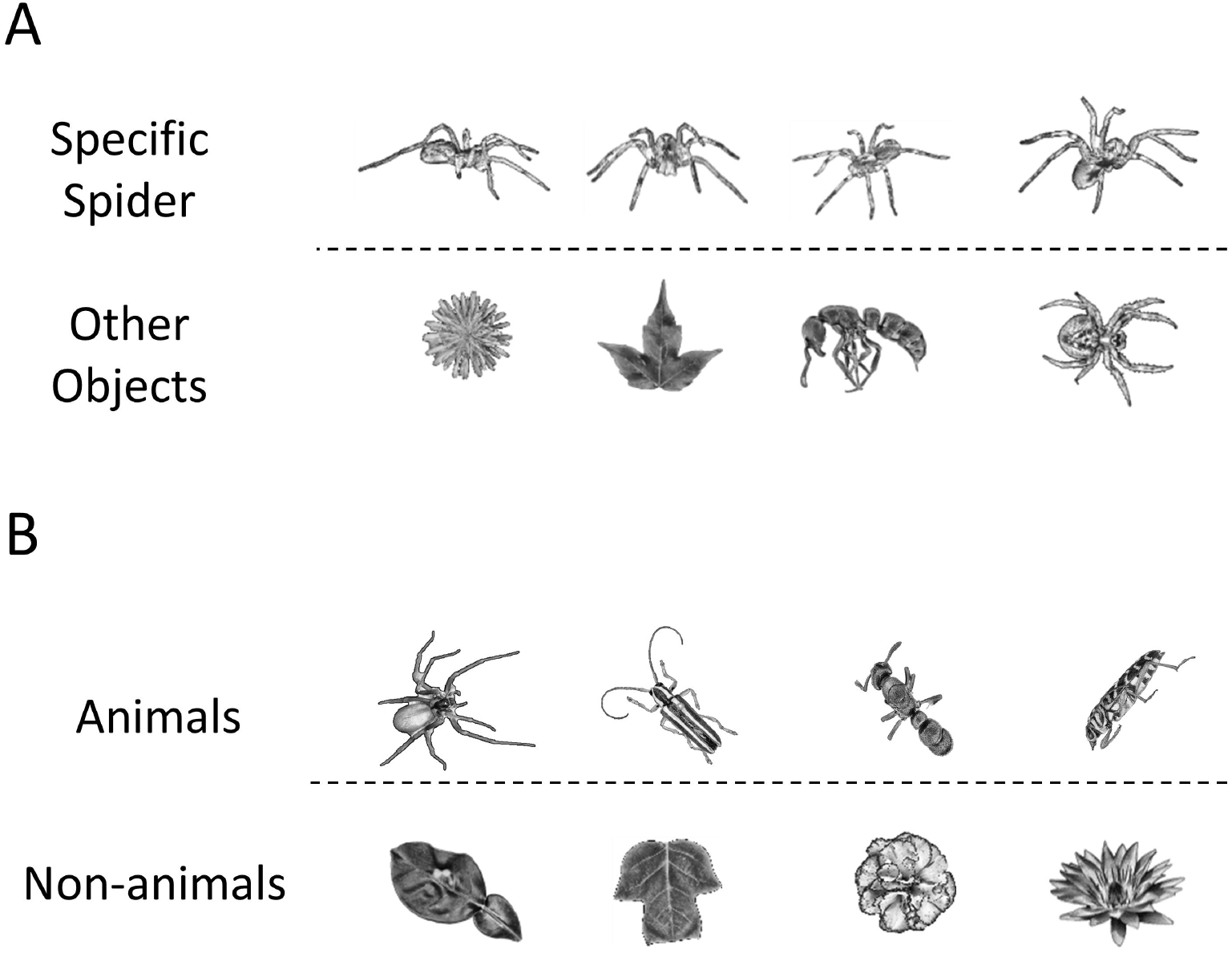
Object recognition problem: Object recognition involves the identification of objects regardless of transformations in size, contrast, orientation or viewing angle. **A.** An example of object recognition of an object, a specific spider in this case, which needs to be identified in the presence of other insects. **B.** An example of object recognition of an object class. In this case, an animal (insect or spider), needs to be recognized in the presence of non-animate objects (leaves or flowers).

Although there is convincing evidence that animals outside of the mammalian clade are capable of object recognition, the mechanisms and formal algorithms underlying this performance remain poorly understood. Pigeons, for example, are capable of categorizing natural objects, human faces, and even emotional expressions (Soto and Wasserman, 2014; Watanabe et al., 2019). Similarly, bees (Avargues-Weber et al., 2010; Giurfa et al., 1997; Werner et al., 2016), wasps (Oliveira et al., 2015; Sheehan and Tibbetts, 2011), and adult zebrafish (May et al., 2016; Oliveira et al., 2015) have all been shown to be capable of conspecific visual identification. A number of studies have indicated that fish also have the capacity to differentiate between different shapes (Mackintosh and Sutherland, 1963; Siebeck et al., 2009), fish faces (Parker et al., 2020), and the archerfish can even be trained to discriminate between human faces (Newport et al., 2016; Newport et al., 2018). Clearly, some of these stimuli, such as human faces, are not ecologically relevant to birds, insects, or fish, nor do we expect fish to possess specific brains areas dedicated to face processing, as is the case for humans. Yet, these findings suggest the existence of a complex visual processing system in the brain that allows for the extraction of the relevant features of an object, its recognition and categorization.

To address these questions concerning the nature of object recognition in non-mammalian vertebrates, we examined the object recognition of natural objects in the archerfish (*Toxotes chatareus*). The rationale for selecting the archerfish draws, in part, on the potential benefits of studying organisms distant from mammals on the evolutionary scale (Karoubi et al., 2016), since this may point to additional visual mechanisms or basic principles. At the same time, the value of the archerfish as an animal model stems from the fact that these fish can be trained to discriminate stimuli visually, even when presented on a computer screen (Ben-Simon et al., 2012; Ben-Simon et al., 2012b; Ben-Tov et al., 2015; Ben-Tov et al., 2018; Gabay et al., 2013; Mokeichev et al., 2010; Newport et al., 2013; Newport et al., 2014; Newport et al., 2015; Newport, 2021; Reichenthal et al., 2019; Schuster et al., 2004; Schuster, 2007; Vasserman et al., 2010). By utilizing this fish’s remarkable ability to shoot down insects and other small animals that settle on the foliage above the water line with a jet of water from the mouth (Lüling, 1963), these fish can be trained to perform an object recognition task and essentially report their decisions using stimuli in the lab. Thus, the archerfish can provide the fish equivalent of a discriminative response by a monkey or by a human when performing a recognition task with a click of a button.

## Methods

### Animals

Eleven archerfish subjects participated in the experiments. Adult fish (6-14 cm in length; 10-18 gm) were purchased from a local supplier. The fish were kept separately in 100-liter aquaria filled with brackish water at 25-29^0^ C on a 12-12 hour light-dark cycle. Fish care and experimental procedures were approved by the Ben-Gurion University of the Negev Institutional Animal Care and Use Committee and were in accordance with the government regulations of the State of Israel.

### Training

After a period of acclimatization, inexperienced fish were gradually trained to shoot at targets presented on a computer screen (VW2245-T, 21.5”, BenQ, Taiwan) situated 35±2 cm above the water level. In the first stage, the fish were trained to shoot at a single black circle on a white background that appeared at random locations on the screen. A blinking black square appeared immediately prior to the display of the target in the middle of the screen and was used as a cue to draw the fish’s attention upward. If the fish shot at the target within 15 seconds from the appearance of the target, it was rewarded with a food pellet. Otherwise, the target disappeared and the next training trial started. The mean response time of the fish ranged from 2 to 10 seconds. The training continued until the fish succeeded in hitting 80% of the targets within 15 seconds.

After the fish learned to shoot at targets on the screen, they were trained to recognize either a specific object or a category through a two-alternative forced choice procedure (Fig. 2A). A session of 20 trials where the fish had to choose and shoot at one of two images was repeated over several (4-10) days to familiarize the fish with the experiment. When the fish achieved a 70% success rate in choosing the designated object or category, it was considered trained and ready for the experiment (examples of the training and experimental procedure for one fish are shown in Fig. 2B). The subsequent experimental trials were recorded and these results were used for the analyses.

**Fig. 2.**
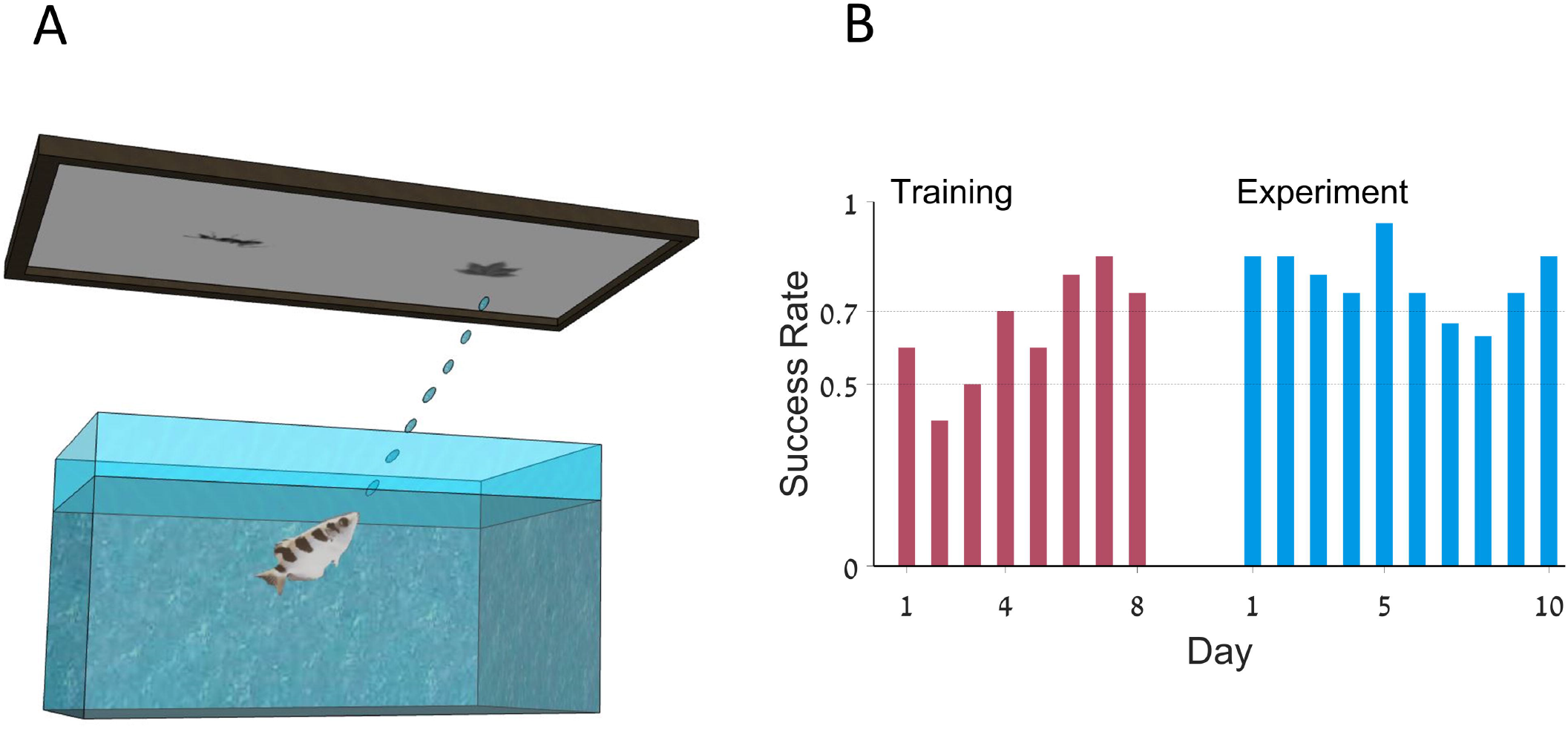
Behavioral experimental setup: **A.** The archerfish is presented with two objects on the screen: a target and a distractor. The fish is rewarded if it selects the target image. **B.** Success rate per day in training process (red) and experiment (blue) of Fish 1.

### Stimuli

For all experiments, we used images of objects familiar to the fish from their natural environment. The images were composed of edible and inedible objects (from the archerfish’s perspective): the inedible objects were either leaves or flowers, whereas the edible objects were either spiders or insects such as a cockroach, an ant, or a beetle. The images of the insects and the spiders were obtained from the BugwoodImages project website (insectimages.org). Images of flowers were taken from the Oxford Flowers 102 dataset (Nilsback and Zisserman, 2008), and images of leaves were taken from the Flavia Plant Leaf Recognition project dataset (Wu et al., 2007). Multiple shots of one specific spider and one ant were taken from the animated 3D models. The models were purchased from the Sketchfub store under standard license (sketchfab.com).

All images were preprocessed using Matlab. All background colors were removed and the objects, after being converted to grayscale, were placed on a white background. The size of the objects was randomized in the following way: the number of the pixels in the image was selected to have a uniform distribution from a discrete set of object sizes. For this purpose, the images were resized to create 5 levels of object area, defined as the number of pixels within the contour of the object: ~10,000 pixels, ~50,000 pixels, 100,000 pixels, 200,000 pixels and 300,000 pixels.

### Experiment 1

We investigated recognition of a specific object in the archerfish. The fish were rewarded with a food pellet if they selected the target. The experiment consisted of ten sessions (each on a different day) with 20 trials per session, and lasted 3 to 5 weeks with 2-3 sessions per week.

There were two types of targets. First, an image of a specific spider was presented to the fish together with a distracting object. The distracting objects were leaves, flowers, insects, or other spiders. The target spider was shown from different viewpoints, with different orientation, size, contrast, and screen locations. All presentation parameters were randomized.

Then, the experiment was repeated with a designated target of a specific ant. The distractors in this case were images of leaves, flowers, insects, or other ants.

### Experiment 2

Whereas Experiment 1 focused on specific objects (and thus on identification), Experiment 2 explored the ability of the archerfish to generalize and categorize various objects into classes. On each trial, two novel images belonging to two different categories – edible and inedible – were presented in random locations on the screen.

### Analysis of image features

To analyze the possible visual features that may help the fish to perform object recognition, we extracted a set of 18 visual features commonly used in image processing from each image (Nixon and Aguado, 2019; Wang et al., 2012; Wen et al., 2009) and then used these features to characterize the images. The following features were used:

1. Object area, defined as the number of pixels within the object’s perimeter. Features describing object compactness:
2. Convex hull area.
3. Convex hull area divided by the object area.
4. Perimeter length.
5. Roundness, defined as the perimeter squared divided by 4π*area. Features describing object curvatures:
6. The number of sharp curves in the object’s perimeter was defined as follows: first, the object’s perimeter was divided into sections with a length of 100 pixels each. Then, a second degree polynomial was fitted to each section, and the polynomial’s second derivative was used as the curvature for every section. Finally, a section with curvature values above the standard deviation of all values for all sections was considered a sharp curve, which yielded the number of sharp curves in the object’s perimeter.
7. Average curvature value of the sections with sharp curves as defined in 6. Features describing object shape eccentricity:
8. Shape eccentricity, defined as the ratio between the foci of the ellipse that surrounds the object and the length of its major axis.
9. Convex hull eccentricity, defined as the ratio between the foci of the ellipse that surrounds the convex hull of an object and the length of its major axis. Features describing object texture:
10. Entropy of light intensities of the objects pixel values.
11. Standard deviation of the object’s pixel values.
12. Skewness, defined as the normalized third central moment of the object’s pixel value distribution.
13. Correlation between the object and a checkerboard: dot product of the object with 5 checkerboards with different checker sizes – 4 to 12 checkers in a row – where the maximum result was used. Other features:
14. Correlation between the object and a star: dot product with a star shape to measure the resemblance of the object to a star.
15. Symmetry, defined as the distance between two halves of an image on the horizontal and vertical axis of the image. All images were rotated to align the major axis of the surrounding ellipse to the x-axis.
16. Symmetry defined as the Euclidean distance between two halves of the image on the horizontal and vertical axis of the image silhouette. Image energy:
17. Mean image energy defined as the average value of all pixels in the image.
18. Total image energy, defined as the sum of all pixel values in the image.

### Support vector machine analysis

We used Matlab Statistics and the Machine Learning toolbox functions to build a Support Vector Machine classifier. The classifier was trained using a matrix with image features and the fish’s responses as labels. The training set consisted of a random 75% of the images. The resulting support vector machine model was tested on the remaining 25% of the images. The labels that the model returned were compared to the images’ true labels and to the fish’s behavioral selection. The average success rate for 20 iterations was used as the model’s success rate. We arranged the features according to the order of their contribution to the model using a greedy algorithm. At every step, the feature that contributed the most to the model’s success was added.

### Statistical analysis

We performed a hierarchical Bayesian analysis to evaluate the behavior of each fish in every experiment. The statistical analysis used R 4.0.4 and JAGS 4.3.0 software to sample the posterior probability distribution of the parameters (Kruschke, 2014). The statistical model we used had a binomial likelihood:

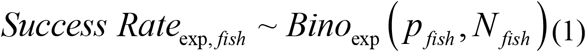

The distribution of the success rate for the different fish, *p_fish_*, was a beta distribution whose mode, *ω*, and concentration, *κ*, were hierarchically determined.

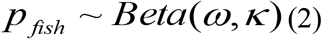

The priors for *ω* and *κ* were chosen to be uniform and very broad.

We used JAGS (Plummer, 2003) to generate 3 chains of 10000 MCMC samples from the joint posterior probability distribution of the *p_fish_* for all fish and experiments. Convergence of the algorithm and sampling properties were tested using the graphical and quantitative methods outlined in Kruschke, 2014. Using the MCMC samples, we calculated the 95% highest density interval (HDI) for the fish’s behavioral success rate, a range of values in which there was a 95% posterior probability of finding the parameter. The success rate of the experiment was considered significantly above the chance level if 95% HDI of its posterior distribution was greater than a region of practical equivalence (ROPE) of 5% around the chance level of 50%. Similarly, if 95% HDI of the difference in success rate between the two experiments included the ROPE of 5% around zero, the success rate was not considered to be different.

## Results

We characterized the archerfish’s ability and processing during an ecologically relevant object recognition task. For this purpose, we conducted two alternative non-forced choice experiments using a continuous reinforcement schedule for correct responses. Generally, the fish was presented with images of natural stimuli that are important in the fish habitat, and was rewarded for an appropriate shooting response. The two stimuli were presented simultaneously on a computer monitor situated above the water tank (Fig. 2A). A shooting response directed at the correct target was rewarded with a food pellet whereas the selection of the other stimulus was not. Successive learning trials were contingent on the fish collecting the reward for the previous correct response. To neutralize the effect of position bias in the fish’s responses, the two targets were presented at a random location on the screen.

### Archerfish can recognize specific objects regardless of differences in contrast, size, and viewing angles

We tested whether the archerfish was capable of object recognition. Three archerfish were trained on a set of pictures of a single spider viewed from different angles in a 3D space and then tested on the same spider viewed from other angles (the generalization set), which were not included in the original set (Fig. 3A). The target spider was presented under different conditions such that size, viewing angle, contrast and location varied from trial to trial. The target spider was presented together with another object that could be a leaf, a flower, an insect, or another spider (Fig. 3A, see Methods).

**Fig. 3.**
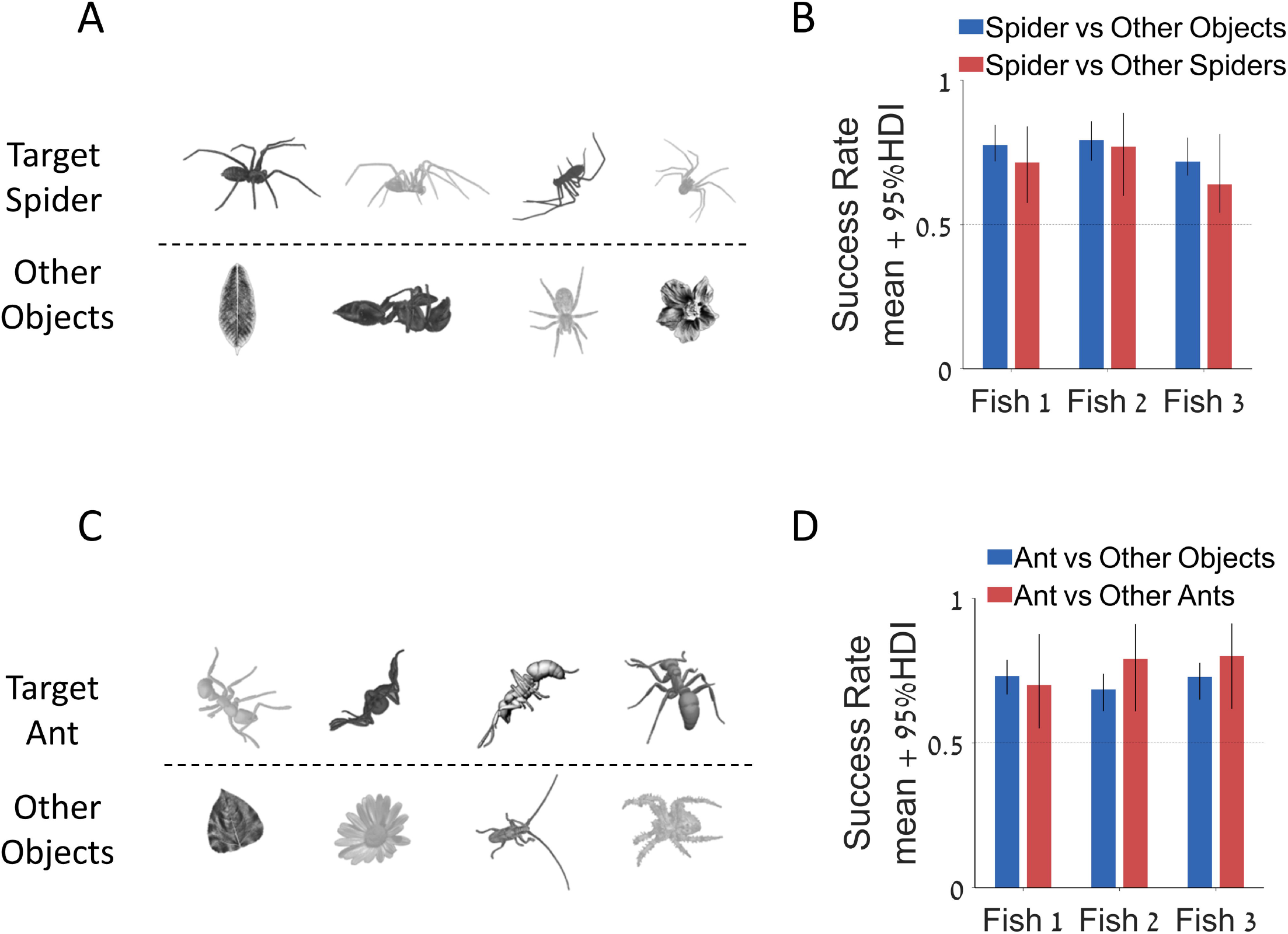
The archerfish is capable of invariant object recognition: **A.** Examples of a single target spider from different viewpoints and with different contrast levels (top row) and other distractor objects and spiders (bottom row). **B.** Success rate of three archerfish in recognizing the target spider: mean + 95% HDI. **C.** Examples of a single target ant from different viewpoints and with different contrast levels (top row) and other distractor objects and ants (bottom row). **D.** Success rate of three archerfish in recognizing the target ant: mean + 95% HDI.

The fish were able to recognize and choose the target spider, both on trials where the second object was not a spider and also against other spiders (Fig. 3B). For all fish in both experiments, there was no overlap of the posterior probability 95% HDI of the success rate with a ROPE of 5% around the chance level (see Methods, Statistical analysis). In addition, the 95% HDI of the difference between the success rate of individual fish on trials with two spiders and the trials with a spider and non-spider completely contained a ROPE of 5% around chance level, indicating that the fish could differentiate the target spider from other types of spiders as well as from other objects.

A similar experiment was conducted with an ant as a target image. The same three fish were retrained to recognize one specific ant that was shown together with other objects, and sometimes with other ants (Fig. 3C). The fish learned to differentiate the target ant from the other objects and also from other ants (Fig. 3D). The success rates in this experiment were not significantly different from the rates in the experiment with a spider target.

### The archerfish can categorize objects into classes and learn to generalize from examples

We tested the ability of the fish to discriminate between the images of two categories of stimuli (Fig. 4A): non-animals (leaves and flowers) and animals (spiders and insects comprising ants, beetles and cockroaches). In this two-alternative-choice-task, in the first stage of the experiment, the animal category was rewarded and the non-animal category was not rewarded. The images were grayscale, normalized to five different sizes, shown at different locations on the screen and were never repeated; that is, each image was used only once (around 1,500 images in total were used in the experiments). After two to eight days, the success rate of the fish reached a plateau that was significantly above chance level (Fig. 4B). The lower boundary of 95% HDI for the fish with the lowest success rate was just above 60%. The higher boundary of 95% HDI for the fish with the highest success rate was above 80%.

**Fig. 4.**
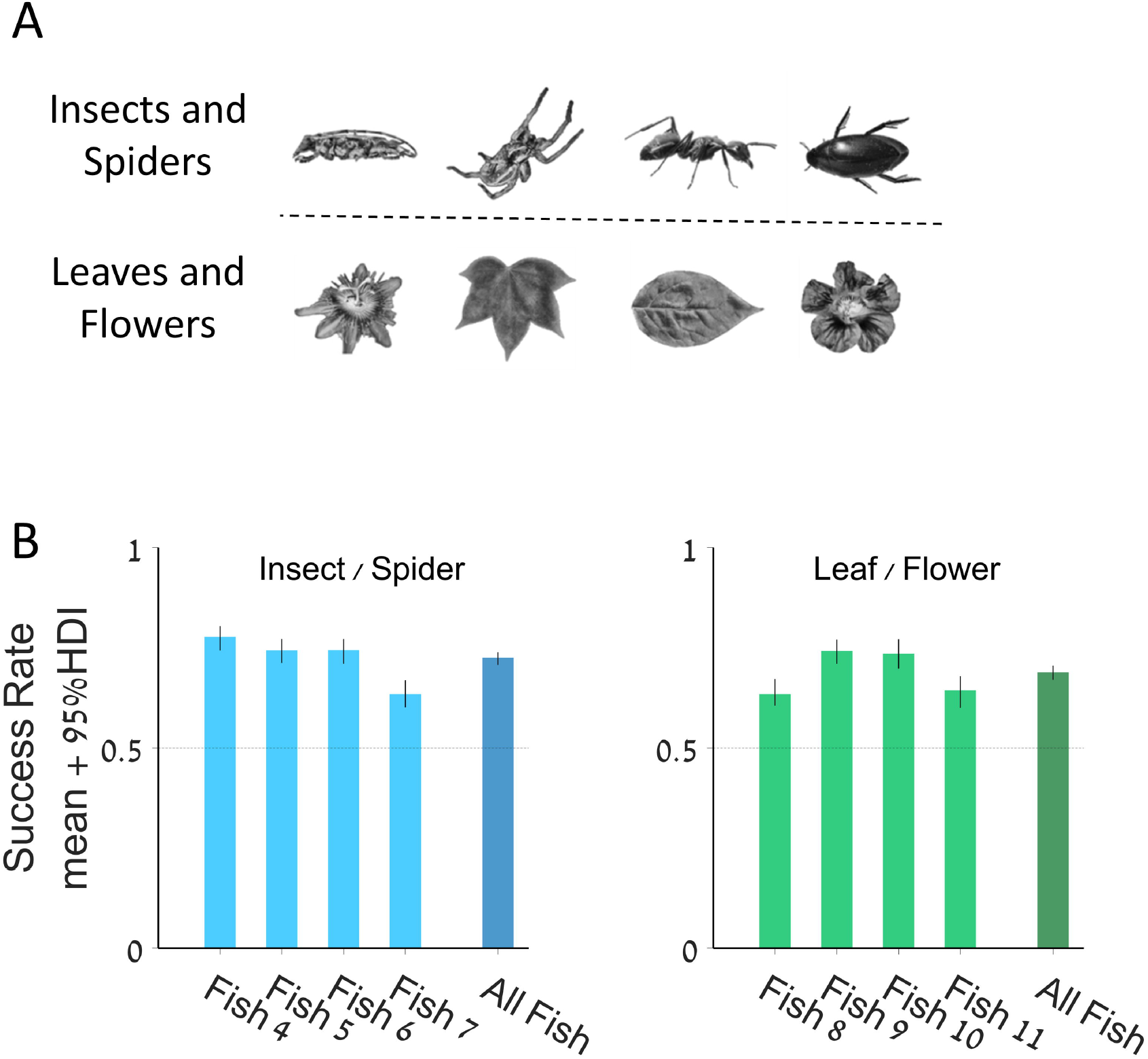
The archerfish can categorize novel objects into groups: The fish were trained to categorize animal and non-animal objects. **A.** Examples of animal objects (insects and spiders, top row) and non-animal objects (leaves and flowers, bottom row). **B.** Success rate of 8 fish in selecting an object from its designated category: mean + 95% HDI. Fish 4 to 7 were rewarded for choosing an animal; fish 8 to 11 were rewarded for choosing a non-animal.

To test whether archerfish are predisposed to shooting at animals rather than plants, we tested four additional fish, which were trained to shoot at the non-animal targets (i.e. non-edible). Again, we found that the archerfish were able to select the non-animal targets at a significantly higher level than chance (Fig. 4B). This is an indication that the archerfish is not hardwired to select an animal.

### The archerfish can use five complex visual features to perform object recognition

To identify the visual features used in the behavioral task, we built a model that simulated the process of object selection in the fish and fit the model to the response data we collected on the fish target selection. The model was composed of two branches of information processing, each processing one stimulus image in parallel (Fig. 5A). Each image recognition module was composed of a feature extraction stage followed by a classifier. The result of the computation by each module was fed into a decision module which, after adding execution noise, led to the behavioral choice of the model. The decision was made by comparing the classification output and verifying its consistency. If the classifications were different, the decision followed suit to the desired class. If both classifiers returned the same category, the decision was made randomly by sampling a Bernoulli distribution with 0.5 probability of success. The behavioral noise itself was added to the decision vector: a decision response was flipped for pairs of images with a probability matched for each fish separately – from 0.65 to 0.8 – to get a success rate fit for the behavioral result.

**Fig. 5.**
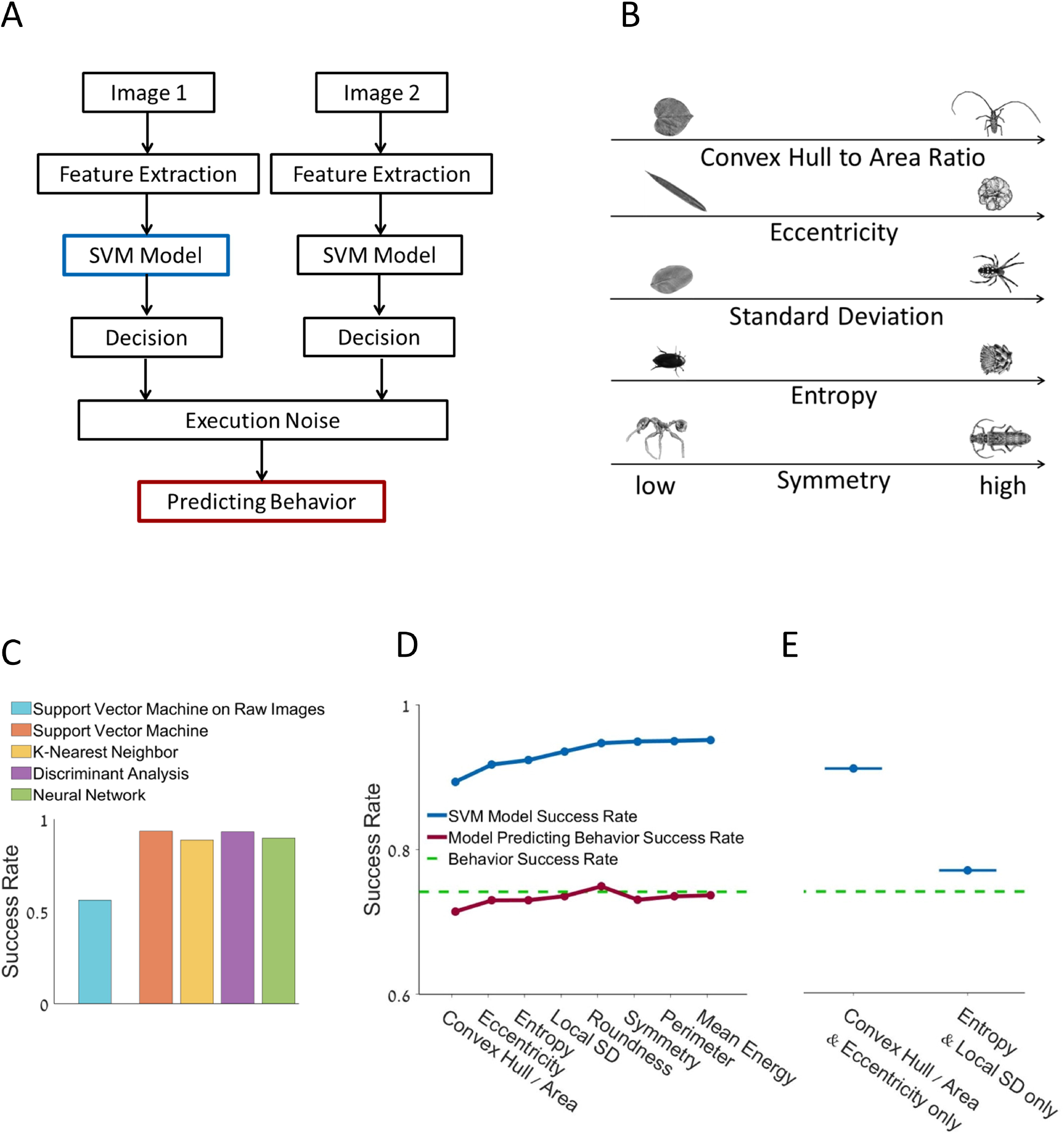
Model building: **A.** In a behavioral experiment, the fish was exposed to two objects, made a decision about the object category and executed a shot. The support vector machine classifier was fed with the features extracted from the images. **B.** Examples of the extracted visual features. **C.** Success rate of different classifiers: support vector machine classifier using raw images, support vector machine classifier using extracted features, k-nearest neighbor, discriminant analysis and neural network. **D.** Support vector machine classifier success rate in predicting the objects’ true category (blue line) and the model’s success rate in predicting fish selection (left, red line). Separate features are added in the order of their contribution to the classifier’s success (left); success rate using only two shape features and two texture features (right). **E.** Support vector machine classifier success rate for combinations of features: two features of shape and two features of texture.

For the classification module we used a support vector machine classifier. The support vector machine was fed by visual features extracted from each image (examples in Fig. 5B, see Methods). We extracted a set of 18 features from each image and then used the support vector machine to build a classifier based on the fish’s responses to the targets and on the extracted features. The features were selected heuristically for the image set (see Methods).

We compared the performance of the support vector machine classifier trained on the raw images to a classifier trained on a feature matrix and found that the use of features significantly improved its performance. We also tried classifiers other than the support vector machine. There was no significant difference in their performance, so we continued with the support vector machine and features for the remainder of the analysis (Fig. 5C).

The classifier was built in an iterative manner, starting with the most informative feature; i.e., the feature with the highest success rate when used in the model separately, then adding the next most informative feature and so on, until the predictive value of the model became saturated. We used a standard training set, verification set and test set to avoid over-fitting the model. Although this was a greedy algorithm that could not guarantee an optimal solution, it still provided a lower bound for the optimal performance.

To test the model (Fig. 5A), we used it to simulate the behavioral experiment. The recognition rate at the output stage of the model matched the behavioral success rate of the fish (Fig. 5D), indicating the capability of the model to capture the statistics of fish behavior. Next, we analyzed the model structure to reveal aspects of the fish’s decisions.

### Shape is more important than texture in the archerfish object recognition

Fig. 5D shows that using only the first five features that describe an object’s shape compactness (ratio of convex hull to area and roundness), shape eccentricity, and texture (entropy and the local standard deviation), the model’s success rate saturated. Using these 5 features, the model achieved a success rate of 94% compared to a success rate of 95% on all 18 features.

We calculated the model’s success rate given only the two first shape features; specifically, the ratio of the convex hull to the area together with eccentricity. The model’s predictions were close to saturation, with a success rate of 92% (Fig. 5E). When given only the two most important texture features, entropy and the local standard deviation, the model’s success rate was only 76%. This suggests that shape was more important than texture in the visual discrimination performed by these fish.

To further test the prediction that shape features were more important than texture, we assessed the ability of the fish to perform object recognition after removing all textures and leaving only the silhouette of the image versus removing all the shape information and leaving only texture (Fig. 6A). The experimental procedure was identical to that used in the original categorization experiment.

**Fig. 6.**
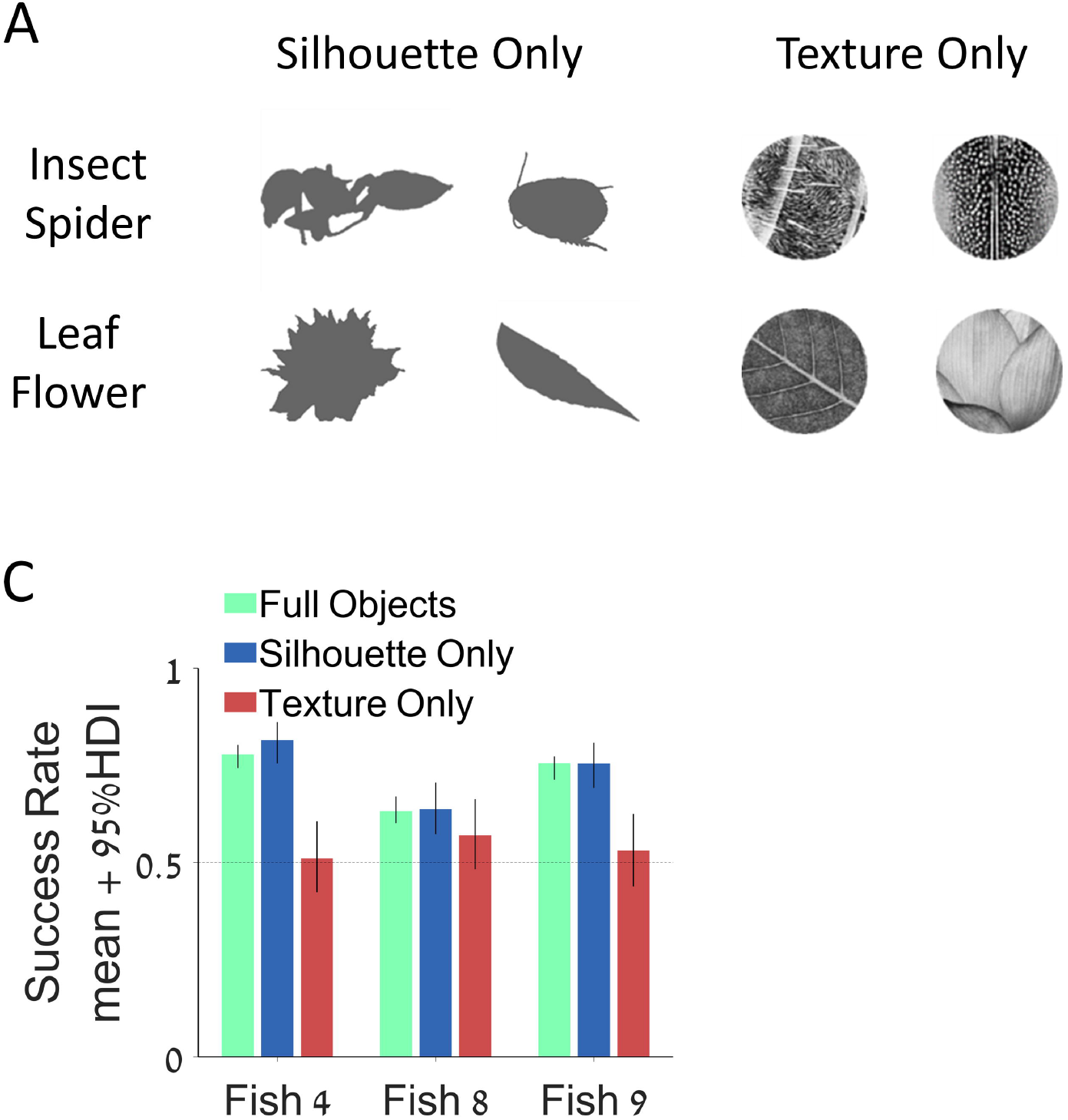
Shape features are more important to recognition than texture: **A.** Examples of animal target silhouettes and textures (top row) and non-animal target silhouettes and textures (bottom row). **B.** Success rate at recognizing the target category in the original experiment with a full object (green bars) and with silhouettes alone (blue bars) and texture alone (red bars) in three fish. No significant difference in the response rate between the original and the silhouette experiments: 95% HDI was above the chance level in all fish. In the texture experiment the 95% HDI range included the chance level of 0.5.

We found that the fish were able to perform object discrimination between animals and foliage when provided only with the shape but failed to do so when provided only with texture (Fig. 6B). This fact, a finding in itself, also increases our confidence in interpreting results from the model.

### Execution noise drives most fish errors

Allowing the support vector machine classifier to learn from the fish behavioral data enabled it to perform this categorization task at nearly perfect performance. Our model (Fig. 5A) attributed the fish errors either to poor classification or to execution noise that was independent of the images. The red line in the graph in Fig. 5D shows the results of adding execution noise to decisions based on the model’s classification. Inspection shows that it matches the performance of the fish closely.

We next conducted a stringent test involving the reexamination of image pairs. It was premised on the assumption that if the internal image processing mechanism has near-perfect performance, most errors are the result of execution noise. We predicted that there would be no significant difference between the success rates of the fish on previously successful and unsuccessful image pairs.

To test this supposition, we repeated the original categorization experiments with four different sets of images: a. the image pairs that the fish identified correctly. b. The image pairs that the fish identified incorrectly. c. The image pairs labelled correctly by the model trained on the results of each specific fish. d. The image pairs that the model labeled incorrectly.

The lower bounds of 95% HDI of the success rate for all fish and all types of targets were well above chance level (Fig. 7A), suggesting that at least part of the errors that the fish made were due to execution noise and not due to the fish object recognition algorithm’s inability to identify an object.

**Fig. 7.**
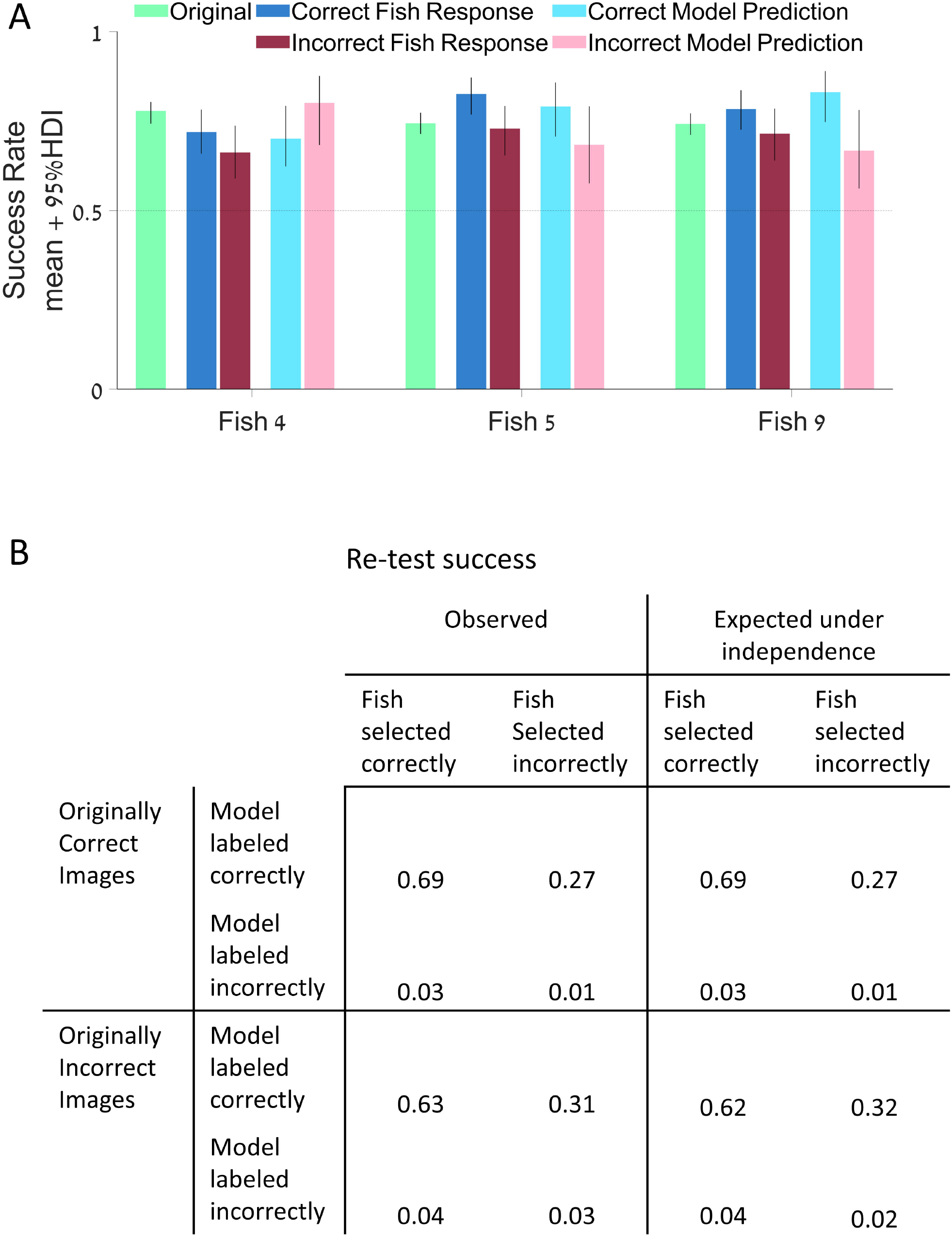
Fish errors are not correlated with object identity: **A.** The original experiment in object categorization was repeated for selected sets of objects: objects that were previously selected correctly by the fish, objects that were selected incorrectly by the fish, objects that the model labeled correctly and objects that the model labeled incorrectly. The 95% HDI of the fish success rate for all sets of objects was above chance level for all three fish that finished all sets. **B.** Portion of images identified correctly and incorrectly by the fish and by the fish-trained model from two datasets: the dataset of images selected correctly in the original categorization task by the fish (top row, left column) and the set of images selected incorrectly by the fish (bottom row, left column); success rate for the same groups expected under independence (right column).

In addition, we compared the selection by the fish and the model for two sets of images: images that the fish identified correctly in the original experiment and the images that the fish identified incorrectly. For each image in the two sets, there were four possible outcomes: both the fish and the model identified it correctly, the fish was correct and the model was incorrect, the fish was incorrect and the model was correct, and both the fish and the model were incorrect. The success rate for all these possibilities did not differ from the success rate expected under independence (Fig. 7B).

## Discussion

Object recognition is an important visual behavior for almost all animals (DiCarlo et al., 2012). However, investigation of the computational aspects of recognition has been confined largely to mammalian species, thus narrowing our understanding of visual processing in general, and limiting the potential for generalizing computational models to new contexts and neural mechanisms. Here, we extended the study of object recognition to a non-mammalian species, to better understand object recognition in general, regardless of the neural substrate or specific ecological context.

In particular, in this work we explored object recognition of natural objects in the archerfish. At its core, visual object recognition binds the stimulus to an internal representation of visual entities that is invariant to most aspects of the stimulus except object identity (DiCarlo and Cox, 2007). This includes invariance to size, contrast, rotation, viewpoint, and illumination, to name only a few, whose variations result in an infinite number of possible projections of the object onto the retina. Our results indicate that the archerfish, like primates and several other species, exhibits this visual function with high accuracy.

Another important and possibly higher level feature of object recognition is the ability to categorize objects by generalizing from examples. We tested this ability in the archerfish by training fish on non-repeating sequences of object images from different classes, and confronting them with novel stimuli that still belonged to the trained classes (as judged by humans). The archerfish were indeed able to generalize across wide range of possible objects and successfully perform the task.

We analyzed fish behavior using a model that aimed to mimic fish behavior. The model was built as a three-stage-cascade composed of visual feature extraction, classification with a learned classifier, and incorporation of additive execution noise before the final decision was made. When we trained the classifier based on the selection made by the fish, we found that it achieved almost perfect performance in predicting the true labels of the objects. Furthermore, it exhibited a hierarchy between features (Fig. 5D), suggesting that the fish attributed more importance to shape feature than to texture features. The model also supported the hypothesis that classification errors were mainly due to execution noise and were not image specific. We tested these two hypotheses with additional experiments and confirmed them both.

### The neural basis of object recognition in the archerfish

Studies suggest that information processing underlying object recognition in the mammalian brain is organized hierarchically and is anatomically located in the ventral stream of the visual cortex (Bracci et al., 2017; Felleman and Van Essen, 1991; Grill-Spector et al., 2001). A visual signal is transferred from the retina to the primary visual cortex V1, where basic features such as oriented lines and edges are extracted (Felleman and Van Essen, 1991; Rust et al., 2005). Information is then transferred through several cortical areas, which select for combinations of visual features, such as orientation and spatial frequency, as well as for higher level geometric features such as curvature (Hegde and Van Essen, 2000). Further downstream, neurons in the inferior temporal cortex have been reported to process complex object features and be tuned to specific object classes such as faces or body parts (Cadieu et al., 2007; Fujita, 2002; Gallant et al., 1996; Lehky and Tanaka, 2016).

Less information is available on visual processing in the archerfish. Previous work on the archerfish have examined visual neural processing in the retina (Segev et al., 2007) and in the optic tectum (Ben-Tov et al., 2015; Ben-Tov et al., 2013), the latter being the largest visual processing area in the archerfish brain (Karoubi et al., 2016). The archerfish optic tectum contains processing stages similar to those found in the early visual system of mammals (Reichenthal et al., 2018). However, it remains unclear whether this area is also the main brain region responsible for object recognition in the archerfish or whether other regions, perhaps within the telencephalon, provide critical functions toward that end.

### Previous studies of object recognition in the archerfish

One of the most seminal studies on object recognition in the archerfish focused on human face recognition (Newport et al., 2016; Newport et al., 2018). The findings showed that archerfish could be trained to recognize human faces in that the fish correctly discriminated one specific face from others, though with apparent difficulty since accuracy decreased markedly on rotated versions of the same face. By contrast, our results show that the fish could identify the same object despite various deformations, including rotation. This could be due to the improved recognition capacity related to the stimuli we used, which were chosen for their ecological relevance (recall that we used insects and foliage).

Other studies have examined the ability of archerfish to recognize simple shapes to test various forms of fish visual behavior, including visual search (Ben-Tov et al., 2015; Reichenthal et al., 2019; Reichenthal et al., 2020), symbol-value association and discrimination (Karoubi et al., 2017) as well as the generalization of the abstract concept of same and different (Newport et al., 2014; Newport et al., 2015). The current study is nevertheless the first to reveal ecologically relevant object recognition in this species.

### Considerations in modelling fish behavior and limitations

To assess the influence of the specific classifier, we tested several other classifiers including k-nearest neighbor, discriminant analysis and neural networks. The success rates of these classifiers were similar to ones observed using the support vector machine both for predicting image true labels and for predicting fish behavior (Fig. 5C). Therefore, the choice of the classifier did not appear to significantly affect the results.

In addition, naïve application of the support vector machine on the raw images, by trying to directly reverse-engineer the image features used by the fish, failed. This is probably due to the high dimensionality of the problem at hand (Afraz et al., 2014). For this reason, we implemented a feature extraction approach followed by the application of the support vector machine, which is the standard approach in the field (Brunelli and Poggio, 1993; Chandra and Bedi, 2018). Finally, it should be noted that our findings do not imply that the neural computations underlying object recognition in the archerfish actually employ an identical or similar algorithm to the one generated by our model.

## Conclusion

We examined the ability of archerfish to recognize ecologically relevant objects. Using a model for the fish selection we showed which visual features were used by the archerfish during visual processing. Future studies should explore whether and how these visual features are represented and used in the neural circuitry responsible for object recognition in the archerfish.

## Acknowledgments

We would like to thank Gustavo Glusman, Hadas Wardi and Jolan Nassir for technical assistance. We gratefully acknowledge financial support from The Israel Science Foundation – FIRST Program (grant no. 281/15), The Israel Science Foundation – FIRST Program (grant no. 555/19), The Israel Science Foundation (grant no. 211/15), The Human Frontiers Science Foundation grant RGP0016/2019, the Frankel Center at the Computer Science Department, and the Helmsley Charitable Trust through the Agricultural, Biological and Cognitive Robotics Initiative of Ben-Gurion University of the Negev.

